# Reference-guided pseudotime inference across species and biological contexts

**DOI:** 10.64898/2026.09.04.749461

**Authors:** Natalie Rittenhouse, Ruth Dannenfelser, Galina N Filippova, Vicky Yao, Xinxian Deng, Christine M. Disteche, Ran Zhang

## Abstract

Cells collected at the same chronological age can vary substantially in biological age due to the heterogeneity in the timing of differentiation, speed of maturation, and degeneration. However, existing pseudotime inference methods either disregard chronological time information, or rely on accurate time-series labels within similar species or biological conditions of interest. As a result, both types of strategies often fail to faithfully order cells from biological contexts without reliable time labels, along the desired axis of interest such as human embryonic development or disease progression. Here, we propose Cavebear, a machine learning framework that enables pseudotime inference in a query species or condition guided by scRNA-seq time-series profiles from a reference species or condition. Cavebear achieves more accurate developmental pseudotime inference than existing methods and provides *in vivo* temporal mapping for *in vitro* experiments. Furthermore, we illustrate the potential of Cavebear to study cellular-level disease progression in human patients using mouse cancer development models as references. By transferring temporal information across species and conditions, Cavebear enables systematic investigation of biological variation in contexts where such annotations were previously unattainable.

## Background

Cellular profiles evolve over time during processes such as development, aging, and disease. Characterizing these temporal changes can reveal the genes and regulatory mechanisms driving cell fate decisions, tissue homeostasis, disease development, and progression. However, individual cells are not synchronized in their time-relevant changes, and cells collected at the same chronological timepoint can vary in pseudotime along the target temporal axis [1,2]. For example, cells within an embryo harbor diverse differentiation and maturation states [2], and cells collected from the same tumor can show distinct stages of malignancy or dedifferentiation [3].

To characterize temporal changes in cellular profiles along a given time axis for scRNA-seq measurements, unsupervised pseudotime inference methods such as Monocle [2,4], diffusion pseudotime [5] and Palantir [6] have been widely used to order cells based on their locations on a cell neighborhood graph. A deep learning method, scTour, utilizes neural ordinary differential equations to infer cellular dynamics across datasets [7]. These methods have proven powerful; however, unsupervised distance metrics may not optimally reflect distance along a particular axis of interest, such as response to viral infection. Recently, supervised methods including Psupertime [8] and Sceptic [9] have been developed to predict pseudotime under the guidance of time labels in time-series datasets, and have demonstrated improved accuracy for cells collected within the same time series. However, these methods require multiple time labels in the study of interest, and assume that batch effects or time-irrelevant biological variations among samples collected across timepoints are minimal, limiting their applicability in real world settings in which comprehensive time-series measurements are unavailable.

These limitations leave several important use cases unaddressed. First, accurate time labels are often unavailable for human embryos, and the same chronological time label may correspond to different biological states across individuals due to heterogeneity in the rates of disease progression. Second, when studying biological variables where samples are difficult to collect, such as in the human brain or the phases of acute disease, individual studies often cover only one or a few timepoints, providing insufficient temporal labels to supervise pseudotime inference. Furthermore, with the growing use of *in vitro* cellular models based on stem cell differentiation, there is increasing interest in anchoring *in vitro* experiments to an *in vivo* timeline to interpret findings in the context of human development, which existing methods struggle to do accurately.

A successful strategy for learning in undercharacterized spaces, is to leverage information learned from well-characterized references. Most recently, TemporalVAE [10] demonstrated the feasibility of reference-guided developmental time prediction. However, it has primarily been evaluated within closely related biological contexts and does not explicitly correct for batch effects or context-specific transcriptional variation. More broadly, reference-guided approaches have enabled cell type annotation [11–14], cell cycle staging [15], and cross-species gene expression prediction [7,16]. These studies suggest that there are conserved aspects of cellular identity that can be shared across species and biological conditions, raising the possibility that pseudotime information may likewise be transferable across diverse biological contexts. Furthermore, unlike studies in human or *in vitro* systems, model organisms can be extensively profiled under controlled *in vivo* conditions, generating dense time-series atlases that comprehensively profile processes such as development and disease progression [17,18]. We therefore reasoned that these datasets could serve as time references, or “rulers”, for anchoring pseudotime inference in contexts where accurate temporal annotation is limited or unavailable.

Here, we propose Cavebear, a reference-guided framework that leverages publicly available high-quality time-series data from a reference to guide pseudotime inference in undercharacterized settings. We demonstrated Cavebear primarily along the developmental timeline, where dense reference atlases with accurate time labels across contexts provide an ideal benchmark for evaluating and performing reference-guided inference. We further show that Cavebear performs well across three diverse biological applications (Table 1). First, using public scRNA-seq time-series from mouse embryogenesis as a reference, we show that Cavebear outperforms existing approaches in predicting developmental pseudotime time in zebrafish across multiple evaluation scenarios. Second, we apply Cavebear to provide an *in vivo* temporal reference for cells from mouse embryoid bodies and human cortical organoids, indicating general agreement of *in vitro* organoid differentiation to *in vivo* states. Finally, using the mouse reference to guide pseudotime inference of matched tumor and adjacent normal tissues from colorectal cancer patients, Cavebear finds heterogeneous cellular-level progression patterns in both tumor and adjacent normal tissues that capture the unique differences in solid and blood tissue types. Together, these evaluations highlight Cavebear’s utility as a transfer learning tool for uncovering mechanistic cell changes across time in contexts that were previously inaccessible.

**Table 1:**
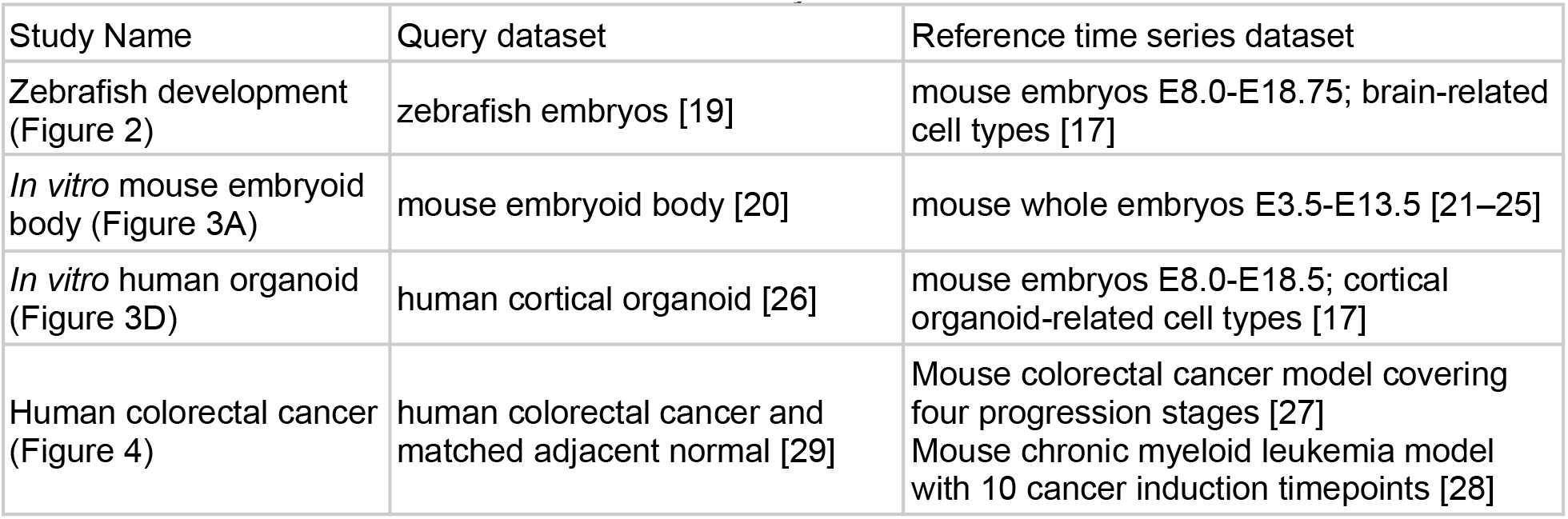
Overview of datasets utilized in this study.

| Study Name | Query dataset | Reference time series dataset |
| --- | --- | --- |
| Zebrafish development (Figure 2) | zebrafish embryos [19] | mouse embryos E8.0-E18.75; brain-related cell types [17] |
| *In vitro* mouse embryoid body (Figure 3A) | mouse embryoid body [20] | mouse whole embryos E3.5-E13.5 [21–25] |
| *In vitro* human organoid (Figure 3D) | human cortical organoid [26] | mouse embryos E8.0-E18.5; cortical organoid-related cell types [17] |
| Human colorectal cancer (Figure 4) | human colorectal cancer and matched adjacent normal [29] | Mouse colorectal cancer model covering four progression stages [27]<br>Mouse chronic myeloid leukemia model with 10 cancer induction timepoints [28] |

## Methods

### Data preprocessing

Cavebear uses raw gene expression count matrices and each cell’s corresponding batch, species, or condition factor as input. For each query study, we chose the reference dataset based on public scRNA-seq time-series profiles that cover corresponding timepoints and cell types relevant to the biological context of interest, as detailed in Table 1. No additional filtering or normalization is performed.

#### Zebrafish development study

We downloaded scRNA-seq time-series measurements from Saunders et al. 2023 and retained wild-type cells annotated to the CNS group, yielding 368,618 cells spanning 37 broad cell types and 18 developmental timepoints from 8h to 96h post-fertilization [19]. For the mouse reference, we downloaded mouse embryonic developmental scRNA-seq data from Qiu et al. 2024 [17]. We first subsampled cells across 67 timepoints spanning E8 to E18.75 to one million cells, then focused on brain development by retaining cells annotated to the following major trajectories: Neuroectoderm_and_glia, Olfactory_sensory_neurons, Oligodendrocytes, CNS_neurons, Intermediate_neuronal_progenitors, Eye_and_other, Neural_crest_PNS_neurons, Neural_crest_PNS_glia, and Ependymal_cells, yielding 403,270 cells. Mouse and zebrafish data were integrated using genes with one-to-one orthology across zebrafish and mouse, resulting in a combined dataset of 771,888 cells and 9,904 genes [16].

#### In vitro mouse embryoid body study

We downloaded *in vitro* cellular profiles of embryoid bodies from Bonora et al. 2021, with collection times at day 0, day 3, day 7, day 11, and differentiated neural progenitor cells (NPCs) [20]. For the mouse *in vivo* reference, we utilized scRNA-seq that spans mouse gastrulation and organogenesis from the Trajectories Of Mouse Embryogenesis (TOME) database [21], which includes datasets from four studies [22–25]. Mouse embryogenesis cells spanned E3.5 to E13.5, with each of the E8.5 to E13.5 timepoints subsampled to the maximum number of cells in the earlier timepoints (16,909 cells). Mouse *in vivo* and *in vitro* data were combined resulting in a combined dataset with 24,324 genes, 10,605 mouse embryoid body cells and 191,883 mouse *in vivo* cells.

#### In vitro human organoid study

We downloaded human cortical organoid data from Glass et al., 2026, and retained high-quality cells by excluding those labeled as stressed or failing quality control (ie. Stressed = True or IVIV_n_QC_passed = False) [26]. Organoids were originally derived from iPSCs from 7 females and 11 males. As the organoid dataset lacks several major brain cell types present in the mouse reference utilized in the zebrafish development study, we filtered the one-million mouse developmental reference to four neural trajectories: Neuroectoderm_and_glia, Oligodendrocytes, CNS_neurons, and Intermediate_neuronal_progenitors. Mouse and human organoid data were then merged based on gene orthologs, yielding a combined dataset of 14,984 genes, 130,673 human cells, and 403,270 mouse cells. Each sample in the human organoid study was treated as a distinct batch following the original study’s convention.

#### Colorectal cancer analysis

The Cavebear model was run separately on a mouse colorectal cancer (CRC) dataset (4 timepoints, 48,117 cells) [27] and mouse chronic myeloid leukemia (CML) dataset (10 timepoints, 310991 cells) [28] reference to make predictions on human CRC target dataset containing 48 patients and 18,2931 cells [29]. For the mouse CRC reference, we assigned the following time labels to the different cancer progression stages: normal: 0, premalignant: 1, malignant (3weeks): 2, malignant (9weeks): 3. For the mouse CML reference, we assigned the timepoints as tumor induction weeks, where timepoint 0 represents the normal stage. To obtain a robust result, predictions from five independent runs with different random seeds were averaged.

### Cavebear framework

Cavebear trains a cross-species pseudotime predictor in three steps: an unsupervised cross-species alignment step (Step 1), followed by a supervised pseudotime prediction step, in which the predictor is trained on reference time labels (Step 2) and subsequently applied to query cells (Step 3).

To first align cells across species in a shared latent space (Step 1), we adopted our previous method, Icebear [16], which learns low-dimensional, species-agnostic cell embeddings from multi-species single-cell data. Briefly speaking, for each cell, the model takes as input the raw gene expression count vector, concatenated with a one-hot encoded batch and context label. The encoder maps this input to a low-dimensional latent representation,, and the decoder reconstructs the input gene expression from the latent representation conditioned on the context and batch labels. By jointly minimizing the reconstruction and KL divergence losses, the cVAE learns a low-dimensional latent cell representation that captures the major sources of biological variation in cellular profiles while removing batch- and context-specific differences. We randomly held out 10% of cells as the validation set, and tuned the learning rate over {0.01, 0.001, 0.0001}. Other hyperparameters are following Icebear’s default. The optimal model is selected based on the highest cross-context Local Inverse Simpson’s Index (LISI) score [30] calculated in the latent space on the validation set. When aligning *in vitro* and *in vivo* systems, Cavebear adopts the optional step in Icebear, which performs adversarial alignment when larger differences between the reference and query remain visible in the latent space after cVAE training. Briefly speaking, during adversarial alignment, we iteratively optimize a discriminator that predicts the context label from each cell’s latent embedding, together with the cVAE, which seeks to minimize the original cVAE loss described above while fooling the discriminator, thereby reducing context-specific information in the latent space. The discriminator weight is treated as a tunable hyperparameter over {1, 2, 5, 10}, and candidate models are selected using the same LISI-based model selection procedure described above.

Given the species-agnostic cell embeddings learned in Step 1, we then train a temporal prediction model using time labels available from the reference species (Step 2). Specifically, Cavebear trains a fully connected multilayer perceptron (MLP) that takes a cell’s latent embedding as input and outputs a single scalar representing its predicted pseudotime. The model is optimized using mean squared error (MSE) loss between the predicted and observed time labels in the reference species. Reference cells were randomly split into training (80%), validation (10%), and test (10%) sets. The best model is selected with lowest validation loss through a grid search over the following hyperparameters: learning rate in {0.1, 0.01, 0.001, 0.0001}, number of hidden layers in {2, 3, 4}, and hidden layer dimension in {50, 100, 200, 400, 800}.

In the final step (Step 3) we apply the trained pseudotime predictor to the query species or condition. The MLP from Step 2 is applied to cells from the query using their latent embeddings from Step 1, producing pseudotime predictions without any time labels in the query context. No additional model training or fine-tuning is performed in this step.

### Comparison with other methods

Because Cavebear does not use any time labels from the query species, we compared it against state-of-the-art unsupervised pseudotime inference methods.

#### Monocle

Monocle [2,4] infers pseudotime by constructing a minimum spanning tree over cells in reduced-dimensional space. Because Monocle 3 produced a large number of infinite pseudotime values, rendering its predictions unreliable for downstream evaluation, we used Monocle 2 for all comparisons and ran Monocle 2 on zebrafish cells. Additionally, the zebrafish CNS dataset contains 368,618 cells and 32,031 genes, which exceeded available memory, so we ran Monocle 2 separately on each broad cell type as annotated in the original study [19]. To ensure reliable downstream evaluation, we filtered cell types to those with at least 10,000 cells, yielding 14 cell types for comparison.

We evaluated Monocle in two aspects: chronological time consistency and within-timepoint heterogeneity (see Model evaluation for details). In the former, Monocle was run on all cells from each cell type jointly, while in the latter, Monocle was run separately on cells from each individual timepoint and cell type. As Monocle is unsupervised and does not inherently reflect the direction of developmental progression, we resolved the directionality of its predictions by orienting the pseudotime axis such that the majority of cells’ inferred pseudotime agrees with their actual collection time order.

#### Palantir and Diffusion Pseudotime

Palantir [6] and Diffusion Pseudotime (DPT) [5] were applied independently to each broad cell type in zebrafish to benchmark against Cavebear. We used the default hyperparameters specified in Palantir’s online tutorial. Counts were normalized and log-transformed, and genes expressing in less than 50 cells were filtered out. We then identified the top 1,500 highly variable genes, performed PCA, and constructed a shared nearest-neighbor graph (50 neighbors). The root cell was defined as the cell with the minimum value along the first diffusion map eigenvector. Palantir was run with 5 diffusion components and 500 waypoints; DPT used the same root cell with 10 diffusion components for consistency. Both methods were evaluated against Cavebear using pairwise accuracy similar to Monocle, with pseudotime sign-swapped per cell type when mean overall accuracy fell below 0.5.

#### ScTour

Following the instructions in the scTour cross-data predictions tutorial [7], we applied the method using the default hyperparameters on the mouse reference to predict pseudotime of zebrafish cells. Gene expressions are corrected for sequencing depth and subjected to log normalization, 1000 highly variable genes are selected, and alpha_z=0.5, alpha_predz=0.5 are used to train the ordinal differential equation model.

#### TemporalVAE

TemporalVAE was trained on the mouse reference following the TemporalVAE internal preprocessing pipeline [10], in which raw counts are corrected for sequencing depth, log-transformed, and z-scored per gene. As the TemporalVAE cross-species prediction interface was not fully implemented at the time of analysis, we adapted the published reproducibility scripts using two sets of recommended hyperparameter configurations (‘supervise_vae_regressionclfdecoder_exp2_toyDataset’ and ‘supervise_vae_regressionclfdecoder_mouse_stereo’). To accommodate the large cell numbers in our dataset, we used a batch size of 1,000. We trained models across 50, 100, and 200 epochs for each configuration. Zebrafish pseudotime predictions were obtained by applying the trained encoder and time prediction head directly to zebrafish cells without retraining, following the cross-species protocol described in the paper. Due to the absence of a published hyperparameter selection protocol for cross-species prediction, we selected the best-performing model based on Spearman rank correlation between predicted pseudotime and actual collection time in zebrafish cells. We note that this selection criterion uses query-species performance information that would not be available in practice, and therefore represents an optimistic estimate of TemporalVAE’s cross-species performance.

#### Upper baseline

Because pseudotime cannot be measured, there is no gold-standard of pseudotime prediction. To establish a practical upper bound, we used the time labels from the query species time-series dataset to train the time predictor, and apply that to predict pseudotime of cells in the query species. For this task, we randomly selected 10% cells as validation, selected the best prediction model based on MSE on the validation set, and predicted pseudotime for all cells. This type of prediction is not practical in reality, as we usually do not have densely measured time series profiles in the query species / study condition. Furthermore, the upper baseline can be influenced by batch effects. Nevertheless, it represents the best possible prediction to serve as a comparison reference for our cross-species pseudotime prediction evaluations.

### Model evaluation

To evaluate pseudotime prediction, because ground-truth pseudotime is not experimentally observable, we used two complementary evaluation strategies.

#### Chronological time consistency

Pairwise accuracy was used to evaluate whether predicted pseudotime preserved the ordering of experimentally measured developmental timepoints. For each pair of sampled timepoints in zebrafish, pairwise accuracy was calculated as the fraction of cell pairs in which the predicted pseudotime agrees with the direction of held-out time labels. Accuracy was quantified separately for each time gap between sampled timepoints. Compared with a single overall correlation coefficient, this evaluation prevents the performance from being dominated by widely separated timepoints, which are generally easier to distinguish than adjacent stages, and offers a more comprehensive picture of pseudotime prediction across different time scales.

#### Within-timepoint heterogeneity

Cavebear was trained using all mouse reference cells together with cells from a single zebrafish timepoint. To evaluate whether Cavebear resolved developmental heterogeneity among zebrafish cells collected at the same timepoint, we quantified the agreement between Cavebear-predicted pseudotime and the upper baseline using Spearman correlation. Cavebear was run separately on each of the four developmental stages (18, 30, 42, and 96 hours post fertilization (hpf)), spanning early, intermediate, and late development, to maximize temporal coverage while minimizing redundancy among adjacent stages.

### Statistical analysis of cancer progression

We restricted our analysis to 10 patients in the CRC query datasets which had matched normal and tumor samples (patients SMC01-SMC10) [31]. We further filtered out 2 cell types, pDC and mast cells, due to low abundance in either the normal or tumor samples, keeping them only in the overall visualization of cells but excluding them in further downstream analyses. Mean predicted pseudotimes were calculated per patient per sample type over cell level values, where we relied on the annotation of cell types provided with the dataset. To show variation across the cells of particular type for a given patient, we estimated an 95% confidence interval using bootstrap resampling with 1,000 resamples. In all comparisons between tumor and matched normal tissue, we used a paired Wilcoxon signed-rank test. To assess whether pseudotime increases with cancer stage, per patient mean pseudotimes were taken for cancer samples only and fine-grained stages (e.g., IIA, IIIA, IIIB) were mapped to an ordinal scale. For each cell type we calculated the Jonckheere-Terpstra test, with the clinfun R package, using the default 5,000 permutations to look for monotonic increases in pseudotimes. All analyses were performed in R (v4.6.0) using ggplot2 and ggpubr for plotting. When reported, all p-values were corrected for multiple hypothesis testing using Benjamini-Hochberg (BH).

## Results

### Cavebear enables reference-guided pseudotime inference across species and experimental systems

Cavebear is a deep learning framework that leverages scRNA-seq time-series data as reference to predict pseudotime for cells collected from another species or condition without well-annotated time labels. With the hypothesis that major biological processes (e.g., differentiation, degeneration) are conserved across biological contexts (e.g., species, experimental systems), Cavebear first tries to align cells from similar cellular state and pseudotime together, and then trains cellular pseudotime based on labels in the reference and apply it to predict pseudotime of the query. Specifically, Cavebear first trains a conditional variational autoencoder (cVAE) using scRNA-seq profiles collected across reference and query to learn context-agnostic cell embeddings that align cells from similar cellular states together (Figure 1). For applications involving substantially different biological contexts, such as *in vivo* and *in vitro* systems, we incorporated an optional adversarial alignment module to further augment cell alignment in the latent space (Methods). Then, leveraging the known time labels from the reference, Cavebear predicts cellular pseudotime by training a multilayer perceptron to predict reference time labels from these embeddings minimizing mean squared error. Although cells from the query species lack time information, because they are mapped to cells with corresponding cell state and pseudotime, the query and reference species share a common latent space in which cells with similar states and pseudotimes are mapped together. The trained time predictor can then be applied to query-species embeddings to generate per-cell pseudotime estimates. Together, Cavebear’s flexible framework allows users to input their query datasets of interest and enables pseudotime inference from different species, experimental systems, and conditions.

**Figure 1:**
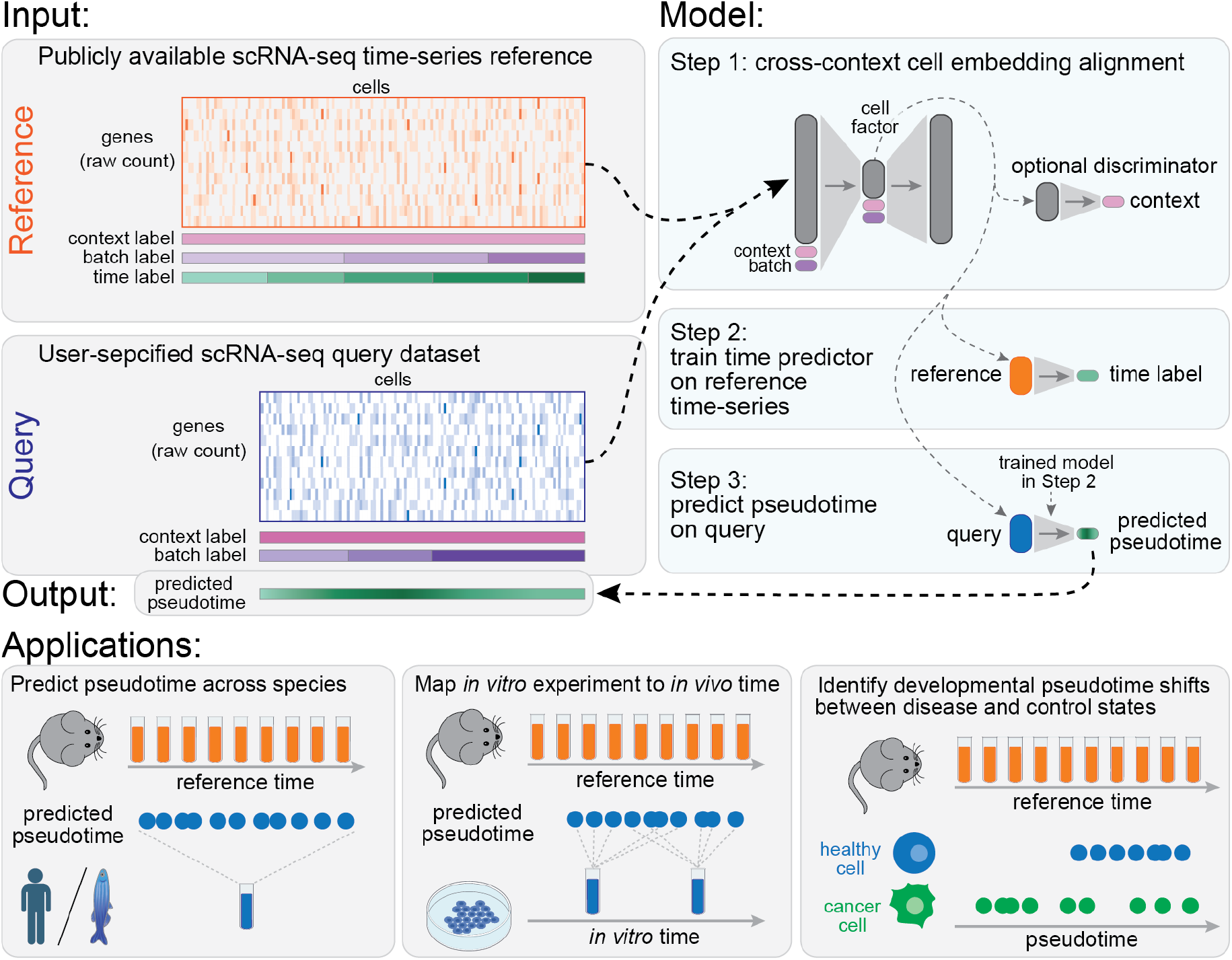
Cavebear’s pseudotime prediction framework. Cavebear leverages publicly available scRNA-seq time-series profiles as a reference to infer pseudotime of cells from a user-specified query dataset. Both datasets are provided to Cavebear, which then adopts a step-wise training framework, where cells are aligned between the reference and query (Step 1), develop a pseudotime predictor (Step 2), and predict cellular pseudotime in the query (Step 3). While Cavebear can have a variety of uses, we demonstrate its application on three separate tasks: predicting pseudotime across species, inferring *in vivo* developmental timing for *in vitro* experiments, and predicting disease progression times for patient cells taken from model systems.

### Cavebear outperforms existing methods on developmental pseudotime inference

To evaluate Cavebear’s capacity at reference-guided pseudotime prediction across species, we first applied and validated Cavebear using scRNA-seq mouse dataset consisting of 403,270 cells from 9 cell types across 67 timepoints and a set of 18 timepoints for 1,222 zebrafish embryos covering 368,618 cells from 33 cell types (Figure 2A, Table 1) [17,19]. Because both datasets consist of whole-embryo profiles with comprehensive time labels, they provide an ideal setting for evaluating Cavebear and comparing it to existing methods. We assigned the mouse as the reference species where time labels are used in model training, and compared the predicted pseudotime for zebrafish cells with known time labels that are unseen from the training process. To enable that pseudotime prediction trained on mouse reference can apply to zebrafish, it is critical that cells having similar identity from the reference and query species are aligned to a species-agnostic space. As a sanity check of the embedding (generated in Step 1), we confirmed that cells from different species align to a shared latent space (Figure 2B). Additionally, the two zebrafish cell types that were not in the mouse reference, hatching gland and notochord, maintained their own distinct identities.

**Figure 2:**
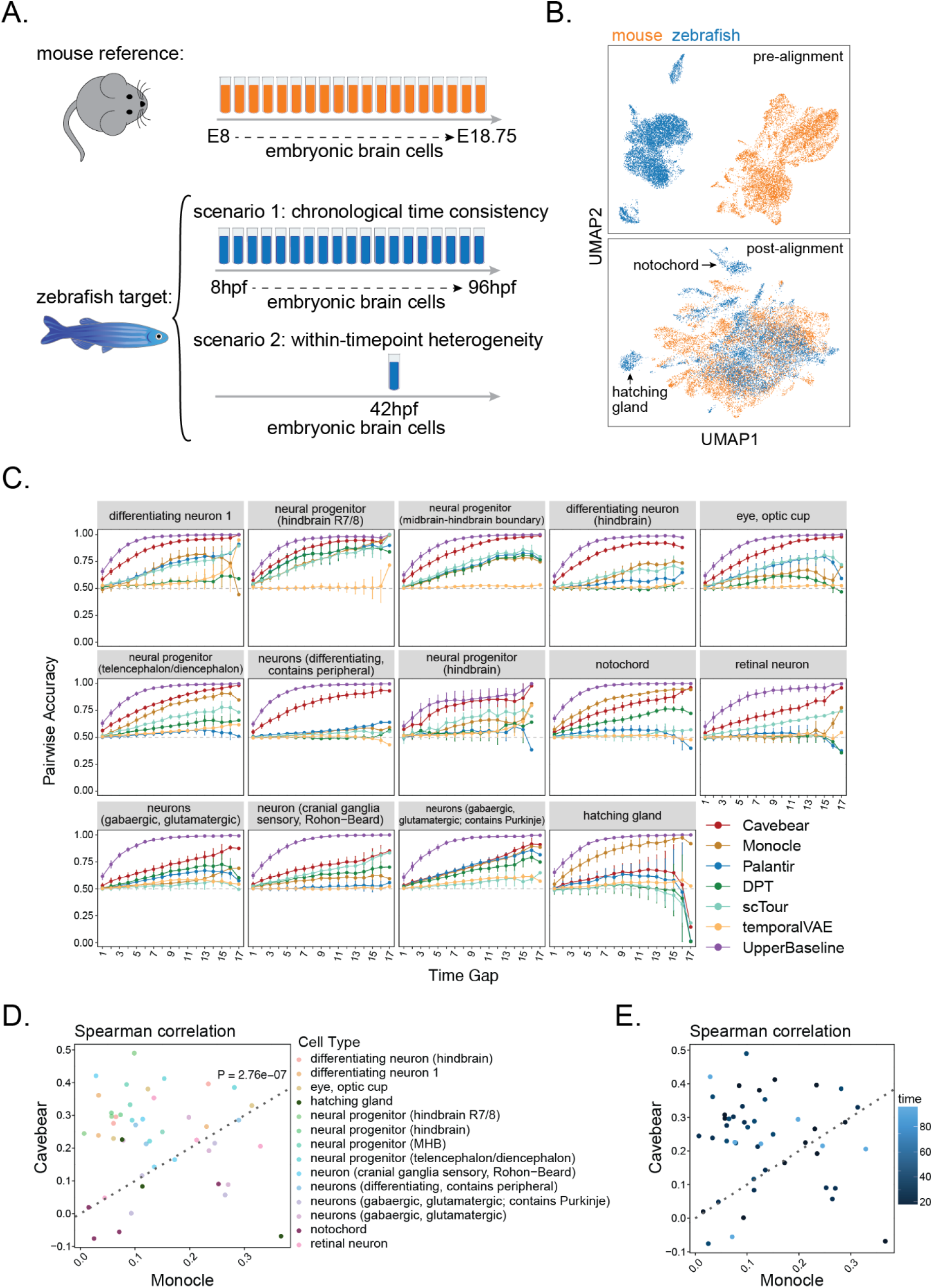
Cavebear outperforms unsupervised pseudotime prediction methods in zebrafish developmental time inference. **(A)** Mouse embryonic brain cells from E8.0 to E18.75 were used as the reference dataset for Cavebear to predict pseudotime in zebrafish embryonic brain cells. For the zebrafish query, two scenarios are tested, a multi-timepoint evaluation where pseudotime is predicted and evaluated across 18 zebrafish samples collected from 8 hours post fertilization (hpf) to 96hpf and a single-timepoint evaluation where predictions are compared within a single collection timepoint. **(B)** UMAP visualization of cell embeddings before and after the initial Cavebear step. Cells are colored and separated by species. The clusters representing zebrafish hatching gland and notochord, which are absent in the mouse reference, are labeled. **(C)** Accuracy of Cavebear predictions in recapitulating the correct temporal order between pairs of cells. For each broad cell type (panel) and each pair of timepoints in the zebrafish query, a pairwise accuracy score is calculated as the fraction of cell pairs where the earlier timepoint receives a smaller predicted pseudotime than the later one. The timepoint pairs are grouped by the temporal gap between timepoints, and dots and error bars represent mean and standard error across all pairs of timepoints, colored by the corresponding prediction method. The dashed line of 0.5 indicates prediction accuracy by chance. Only cell types with at least 100 cells per timepoint are included, and cell types are ordered based on the mean pairwise accuracy of Cavebear across all temporal gaps. **(D-E)** Spearman correlation between predicted pseudotime from Cavebear or Monocle 2 and the upper baseline, evaluated per cell type and collection timepoint. Each dot represents one cell type-timepoint combination, colored by cell type (D) or zebrafish collection time in hpf (E).

Steps 2 and 3 of Cavebear involve training a time predictor on reference timepoint labels and applying it to predict pseudotime in the query dataset. To evaluate this crucial piece of the algorithm, we want to check both the general trends of pseudotime predictions across cell types as well as the heterogeneity of individual cell trajectories. More specifically, because the true developmental pseudotime of an individual cell cannot be directly observed, we rely on the expectation that cells collected at a later developmental timepoint should, on average, be assigned larger pseudotime values than cells from an earlier timepoint. For each broad cell type defined in the original zebrafish query study [19], we therefore tested whether cells from later timepoints were assigned larger pseudotimes than cells from earlier ones. However, given the heterogeneity of developmental pseudotime among cells, some cells from an earlier timepoint may be more advanced than a small fraction of cells from a later timepoint. To calibrate performance accordingly, we constructed an additional “upper baseline” in which pseudotime was predicted by supervised training directly on zebrafish time labels (Methods, Figure 2C). Aggregating accuracy across timepoint pairs, the upper baseline improves as the time gap between timepoints grows. This is consistent with the biological expectation that cells from neighboring timepoints may have overlapping pseudotime due to unsynchronized development, while cells from distant timepoints are unlikely to be at similar developmental stages.

Having established an upper bound for pseudotime prediction performance, we evaluated Cavebear’s pseudotime prediction, which has only seen the time labels in the mouse reference, against the actual zebrafish labels. Cavebear correctly reflects the pseudotime ordering in zebrafish in most pairs of timepoints, and shows increased performance as time gap increases (Figure 2C). Among different cell types tested, Cavebear can better differentiate cellular pseudotime among differentiating and progenitor cell types. Cavebear’s performance is close to the unrealistic upper baseline in hindbrain neural progenitors. These results demonstrate that a reference time-series from the source species can guide temporal inference in a query species with no available time labels.

We further compared Cavebear against existing methods that can predict developmental pseudotime on zebrafish without time annotations. Among them, temporalVAE [10] is the recently proposed direct competitor, as it also uses time labels in the reference to guide pseudotime prediction of the query. Because unsupervised methods – including Monocle [2,4], DPT [5], Palantir [6], and scTour [7] – cannot tell the direction of development, we helped orient their inferred pseudotime to agree with the observed time in the zebrafish query, which is not feasible in practice. Given memory constraints, Monocle, DPT, and Palantir were applied separately to each broad cell type, while scTour and temporalVAE were run across all cell types. Although Cavebear does not use cell type information or query time labels to direct the prediction, it outperforms all competing methods in 12 out of 14 cell types tested, with the trend consistent across timepoint pairs and time gaps (Figure 2C). Cavebear is outperformed by Monocle in hatching gland and notochord, which are not in the mouse reference, despite still achieving the second highest performance. Moreover, Cavebear significantly outperforms temporalVAE across all cell types, which is trained using the same data and labels.

To further investigate the effect of reference choice to cross-species learning, we focused on cell types with cross-species correspondence, and constructed three subsampled references from the original mouse reference by retaining every 4th, 16th, and 64th timepoint, which reduced the number of reference timepoints from the original 67 to 17, 5, and 2. We then trained a separate Cavebear model on each subsampled reference, and compared the three models’ performance against the gold standard time labels in zebrafish stratified by cell type (Figure S1A). Reducing the timepoints slightly decreases performance from 67 to 17 (0.0386 mean accuracy difference), and 67 to 5 (0.0884 mean accuracy difference) with even as few as 2 timepoints still capturing meaningful signal (0.1383 mean accuracy difference). Furthermore, the smallest subsampled Cavebear covering 2 timepoints outperforms Monocle, the best performing previous method, in 6 out of 12 cell types, and outperforms Monocle in 11 out of the 12 cell types with the 17 timepoint model, suggesting that Cavebear is robust to the overall timepoint density of the reference.

Beyond timepoint density, we also asked how the time range of reference affects overall prediction stability. To do that, we divided the original reference to three non-overlapping intervals: early (covering 35 timepoints from E8-E10.75), middle (E11-E14.75 with 16 timepoints), and late (E15-E18.75 with 16 timepoints) and retrained Cavebear (Figure S1B). Interestingly, while the full Cavebear generally performed the best (9 out of 12 cell types), the middle development Cavebear model had slightly increased performance for gabaergic neuron types, likely honing in on the peak of neurogenesis processes which are concentrated during this time period [32]. Meanwhile, the late Cavebear model is the worst performer, struggling to predict pseudotime for cell types such as neuron progenitor hindbrain R7/8, which occurs earlier in mouse embryonic development [17]. References should therefore be selected thoughtfully to cover ranges that are relevant for cells in the target for optimal performance.

**Figure S1:**
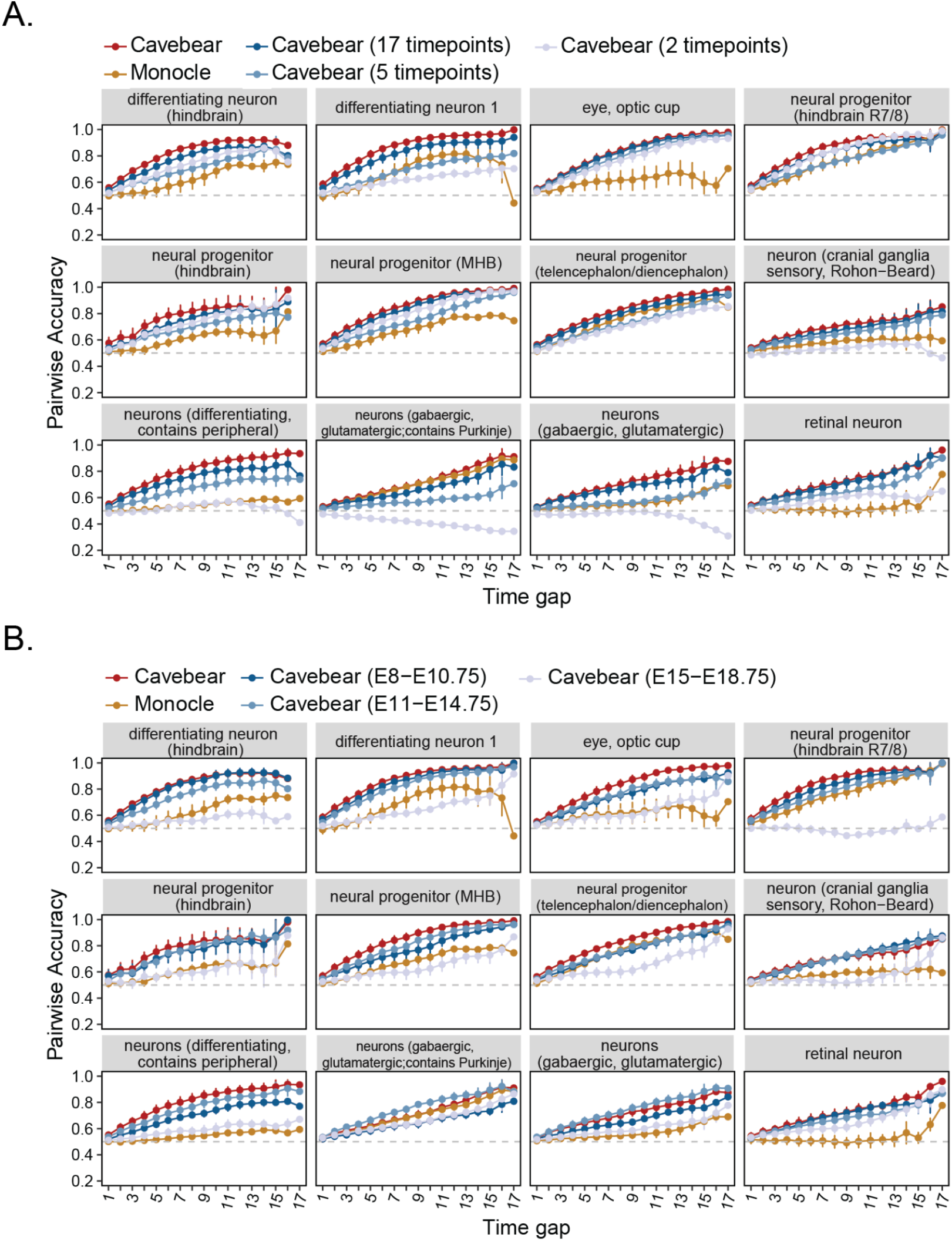
Cavebear’s performance is affected by timepoint density and coverage. **(A)** Accuracy of Cavebear predictions in recapitulating temporal order between pairs of cells. Each color represents a prediction method. Cavebear represents the full model trained on all 67 timepoints in reference, and methods colored in blue show subsetted Cavebear models covering a reduced number of timepoints. Monocle is shown as a comparison baseline. **(B)** Similar to A, the Cavebear model is trained on reference timepoints within three time periods, colored in different shades of blue, to capture how changing reference periods affect results.

### Cavebear disentangles developmental time heterogeneity within a single timepoint

Having validated Cavebear on cells spanning multiple timepoints in the query species, we next asked whether it can resolve pseudotime heterogeneity among cells within a single cell type collected at a single timepoint. This substantially broadens Cavebear’s applicability, as many scRNA-seq studies profile cells from only one timepoint, and cells collected at the same chronological time can vary at different maturation rates along the developmental axis. To evaluate this, we trained separate Cavebear models using the full mouse time-series reference together with zebrafish cells from each of four timepoints (18h, 30h, 42h, and 96h), then assessed predicted pseudotime against the upper baseline per timepoint and broad cell type (Figure 2A). Despite seeing only a single zebrafish timepoint during training, Cavebear’s pseudotime prediction shows positive Spearman correlation with the upper baseline across the majority of cases. Because Monocle’s pseudotime prediction is not oriented, it has equal likelihood of having positive or negative correlation with developmental time. To evaluate Cavebear’s ability on correctly order cells instead of merely being able to tell the direction of development, we gave Monocle an unfair advantage by taking the absolute value of its Spearman correlation with the upper baseline, so that its pseudotime is oriented in the direction that agrees with observed developmental time. This is unrealistic in practice, as no additional time labels or upper baseline is accessible in this scenario. Despite the big advantage given to Monocle, Cavebear prediction shows significantly higher Spearman correlation with the upper baseline compared to Monocle (Figure 2D-E, one-sided paired Wilcoxon signed-rank test). The trajectories where Cavebear underperforms are enriched in those not shared between mouse and zebrafish, consistent with our earlier finding that cross-species transfer is most effective when biological trajectories are conserved between the reference and query. These results suggest that Cavebear can enhance a sample’s single snapshot with meaningful pseudotimes by leveraging reference time-series data from a related species

### Cavebear predicts in vivo timing for in vitro systems

*In vitro* systems are invaluable for studying development and disease when direct access to *in vivo* tissues is limited or impossible, yet mapping and assessing the system relative to *in vivo* timeline remains a major challenge. Motivated by this limitation, we first tested Cavebear on the feasibility of inferring *in vivo* developmental pseudotime for embryoid body (EB) cells. To do that, we used mouse time-series reference data [21] collected from four separate studies [22–25] that covers 19 timepoints from embryonic developmental day 3.5 to day 13.5 to guide pseudotime inference among spontaneously differentiating mouse ESCs collected from day 0, 3, 7, and 11 (d0, d3, d7, and d11) EBs and further differentiated neural progenitor cells (NPCs) (Figure 3A) [20]. Because of the difference of *in vivo* and *in vitro* systems, we incorporated a discriminator to further align *in vivo* and *in vitro* cells to bridge transcriptional gaps between experimental systems. Predicted pseudotime increased monotonically across the five *in vitro* collection timepoints, demonstrating that Cavebear successfully maps *in vitro* differentiation onto the *in vivo* developmental timeline. (Figure 3B).

**Figure 3:**
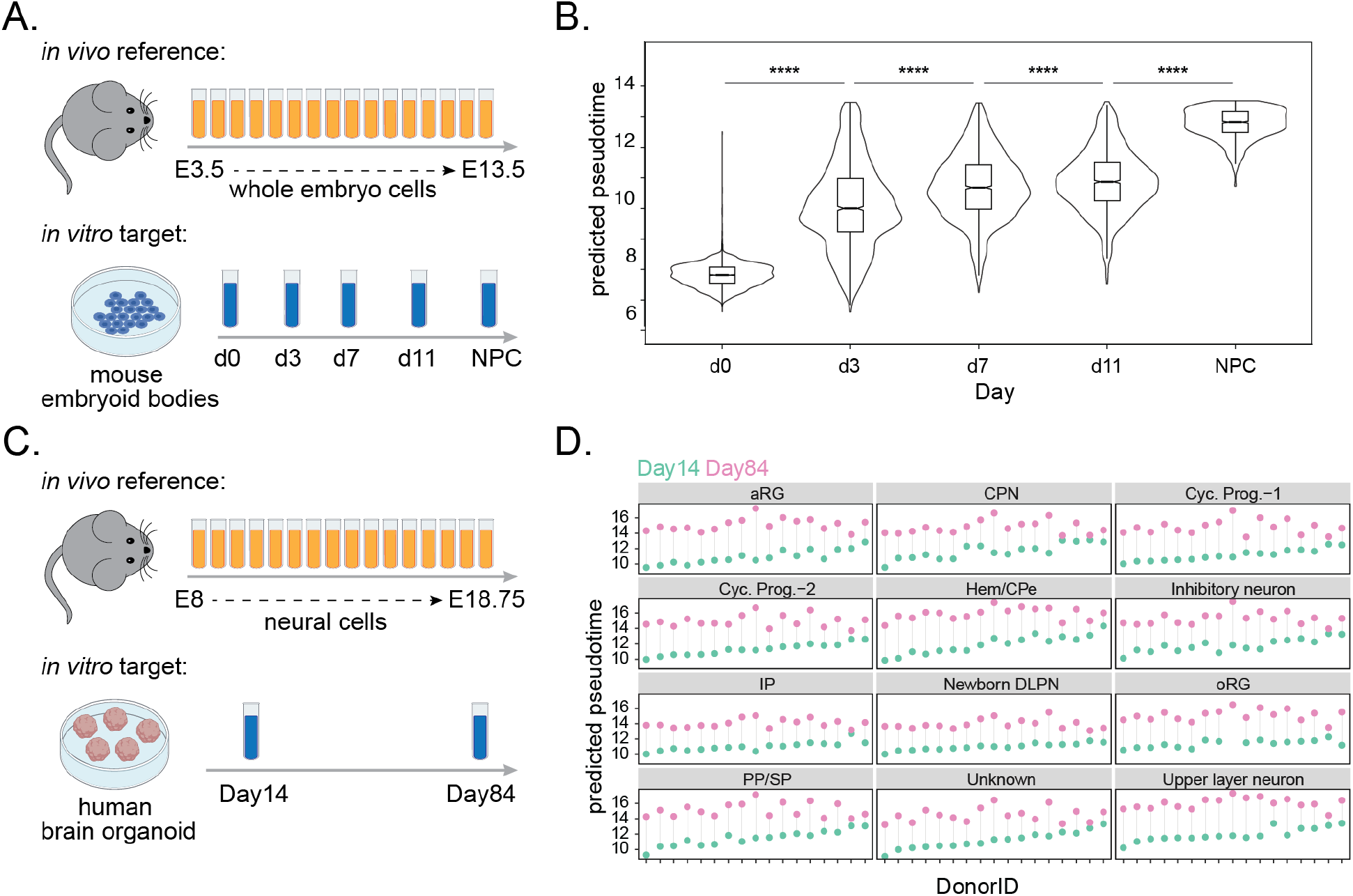
Cavebear predicts *in vivo* pseudotime across *in vitro* experimental systems. **(A)** Mouse embryonic cells from E3.5 to E13.5 were used as the reference dataset for Cavebear to predict pseudotime of *in vitro mouse* embryoid body cells collected at day 0, 3, 7, 11 and as differentiated NPCs. **(B)** Violin plots show the distribution of *in vivo* predicted pseudotime (x-axis) at *in vitro* embryoid body collection times (y-axis). Significance determined by Wilcoxon Rank Sum Test, ****p-adj < 0.0001. **(C)** Mouse embryonic cells from E8 to E18.75 were used as the reference dataset for Cavebear to predict pseudotime of *in vitro* human organoid cells collected at day 14 and 84. **(D)** Differences of median developmental pseudotime (y-axis) between Day 14 and Day 84 organoids derived within the same donor (x-axis), calculated per cell type (panels). aRG: apical radial glia; CPN: cortical projection neuron; Cyc. Prog-1: Cycling Progenitor cells 1; Cyc. Prog-2: Cycling Progenitor cells 2; Hem/CPe: hem/choroid plexus; IP: intermediate progenitor; Newborn DLPN: deep layer projection neuron; oRG: outer radial glia; PP/SP: preplate/subplate.

In addition, we applied Cavebear to predict *in vivo* developmental pseudotime for human cortical organoids [17,26]. Using the mouse reference to guide pseudotime inference of organoids collected at Day 14 and Day 84, we found that Day 14 cells were assigned earlier pseudotime than Day 84 cells across all cell types and donors (Figure 3D) suggesting that our framework is robust across confounding factors. We flagged one donor, 429591, which shows a delayed developmental process comparing Day 84 with Day 14 (Figure 3D, second donor to the right). Interestingly, Donor 429591 is a male donor that shows the smallest cortical surface area at 6 months of age and is the second to last among all donors (male and female) [26]. Because male donors tend to have larger brain regions than females [33], the small surface area suggests there might be neurodevelopmental delay in the donor, as supported by the consistent delay of pseudotime across cell types in Day 84. In addition, mapping organoid pseudotime onto the mouse *in vivo* reference suggests that Day 14 and Day 84 organoid cells most closely resemble mouse embryonic stages around E10 and E15, which is expected as neurogenesis is occurring in both model systems during these time periods. These results demonstrate that Cavebear provides an approximate developmental coordinate system for evaluating and interpreting *in vitro* observations in the context of *in vivo* development.

### Cavebear reveals gradual cancer progression patterns in colorectal cancer at the cellular level

To illustrate Cavebear’s utility beyond development, we applied it on a human colorectal cancer (CRC) dataset with mouse disease model references to reveal disease progression patterns at cellular resolution. Such a view is not typically accessible since patient samples for most diseases are limited, typically to a singular timepoint, whereas model systems can capture and monitor the development and progression of disease across multiple timepoints. Specifically, here, we used a target CRC human dataset [29] with 10 matched normal and tumor samples with an inducible genetic colorectal cancer mouse model reference covering normal, premalignant, malignant at 3 weeks and malignant at 12 weeks [27]. In the reference, each phase is represented as stepwise timepoints ranging from 0-3, thus we expect cells that are normal to have low predicted pseudotimes (<1), with more aggressive / transformed cells to have high pseudotimes (>=1.5). Plotting the individual cells across the 10 patients, we generally see higher pseudotimes in the cancer samples relative to the matched adjacent normals, with the majority of cells across cell types having pseudotimes > 1 in the tumor samples (Figure 4A). Interestingly, we also observe a subset of cells with malignancy-associated pseudotimes (>1) in the matched normals, scattered predominantly in the blood associated cell types, T cell, NK cell, B cell, plasma cell, macrophage, and dendritic cells, suggesting that some malignancy-associated changes persist in the blood of adjacent normal tissue. This dichotomy is particularly apparent when looking at the average pseudotimes per patient and per cell type (Figure 4B), where the three solid tissue-associated cell types, endothelial, epithelial, and fibroblast, have significantly increased (paired Wilcoxon rank sum test, P adj. < 0.001) pseudotimes in the tumor relative to their matched normal sample. Further analysis revealed that the patient (SMC03) with the highest overall average pseudotimes in the tumor solid tissues (fibroblast = 1.98, endothelial=1.71, epithelial=1.66), has the most mutations and the most advanced stage of the 10 patient set. More broadly, we looked for patterns of progression across patients by checking for associations between patient pseudotimes in the tumor and advancing tumor stage (Figure 4C). While not all stages were represented across patients, we see general trends with increasing average pseudotimes across cell types, with the association reaching significance in 2 out of the 3 solid tissues (fibroblast and endothelial). Taken together, these findings suggest that Cavebear inferred pseudotimes are effectively leveraging progression related signals from the mouse reference.

**Figure 4:**
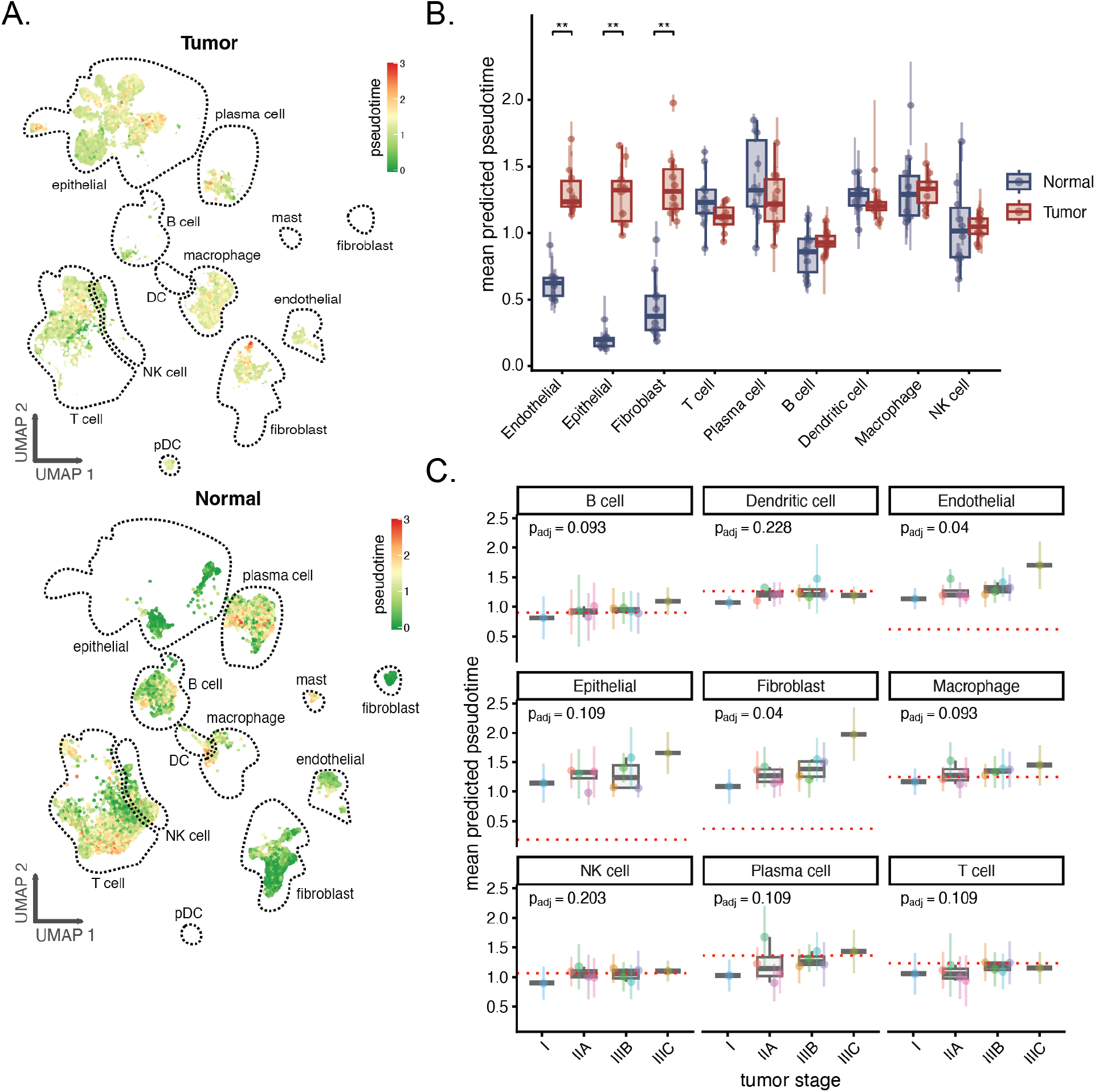
CRC progression at the cellular level revealed by Cavebear. **(A)** UMAP plots showing predicted pseudotimes for cells taken from 10 CRC patients. Cells exclusive to tumor samples (top) have overall higher pseudotimes than those taken from the matched normal samples (bottom). **(B)** Average per patient pseudotimes per cell type are visualized as dots with lines extending to highlight a 95% CI. The average over all patients is shown in the overlaid boxplot with colors corresponding to the sample type (normal in blue and tumor in red). Stars above indicate Benjamini-Hochberg adjusted p-values at different levels of significance (** p<0.01, *p<0.05). **(C)** Average patient pseudotimes for tumor samples only along with their corresponding 95% CIs are again plotted as colored dots, with a boxplot summarizing the average pseudotimes per cell type across patients for stage groups. BH adjusted p-values are calculated for each cell type using the Jonckheere test.

Given that not all diseases have corresponding *in vivo* or *in vitro* models measuring progression profiles, we further asked whether Cavebear can extract progression signals that generalize across a related but distant disease, such as a pair of differing cancer types. To test this, we retrained Cavebear for the human CRC query dataset using a chronic myeloid leukemia (CML) progression mouse model covering 10 timepoints, normalized from 0 (pre-malignant) to 1 (advanced disease) as the reference [28]. Although several cell types are not present in the reference and progression is represented on a different time scale, two solid tissue types, endothelial and fibroblast, still showed a significant increase (paired Wilcoxon rank sum test, P adj. < 0.05) in progression pseudotimes between the paired tumor and adjacent normals (Supplementary Figure 2). Furthermore, the heavily mutated patient, SMC03, again showed the highest average pseudotimes for both of these solid tissues. Despite using a blood cancer reference, which may have substantially different underlying disease machinery from solid tumors [34], Cavebear was still able identify on common cellular features associated with progression to find consistent signal across cancer references.

**Supplementary Figure S2:**
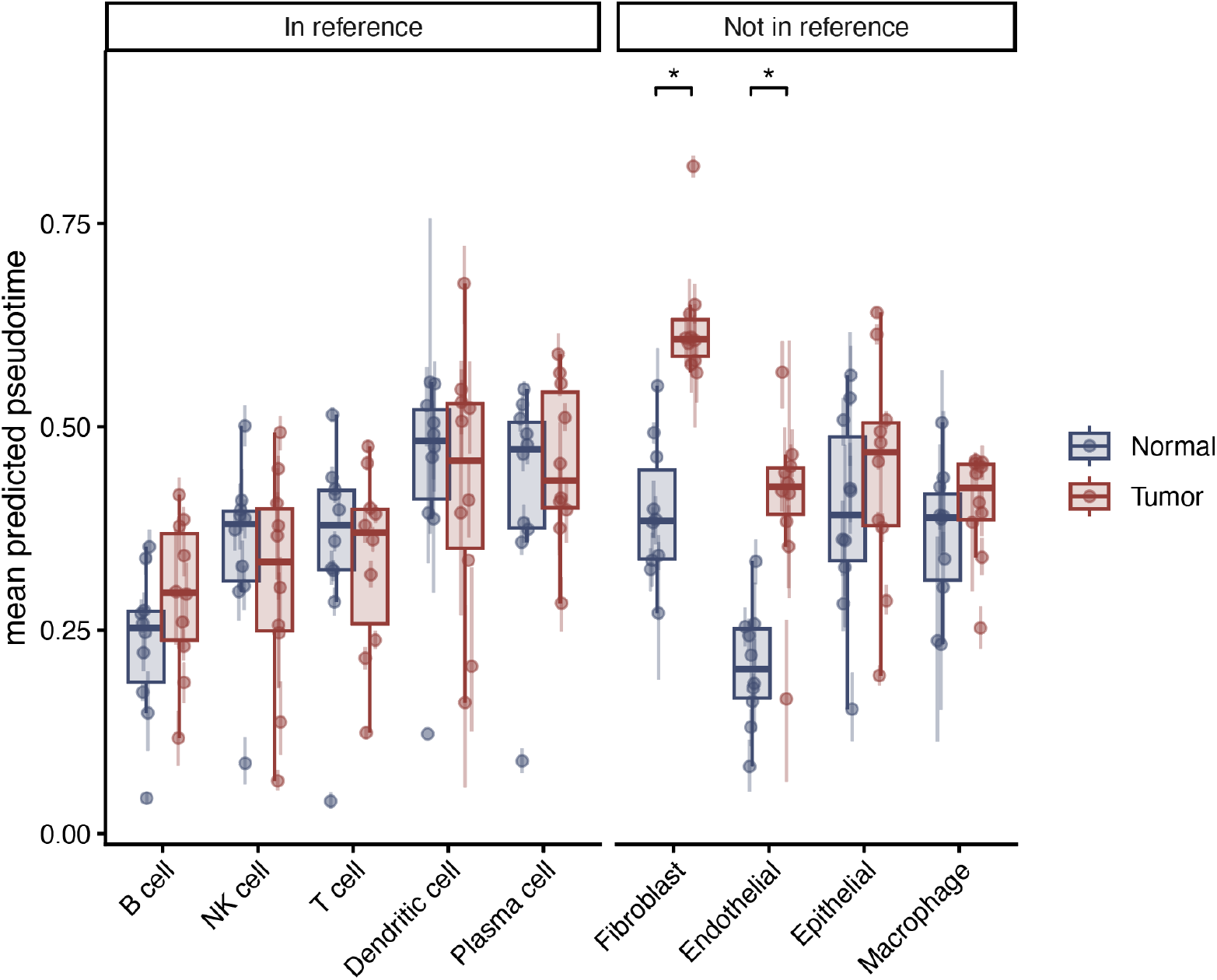
Cancer progression signals in a human colorectal target dataset can also be detected when using a CML mouse reference. (indicates adjusted p-value at different levels of significance: ** < 0.01, * <= 0.05)

## Discussion

Pseudotime inference on single cells provides valuable information about continuous cellular changes underlying biological processes. However, current pseudotime estimation methods either lack a biology-informed reference or are limited to contexts where accurate time labels are available. In this study, we demonstrate that temporal information can be transferred from well-characterized reference systems to different species, experimental models, and across diseases, enabling the reconstruction of biological pseudotime in contexts where direct temporal annotations are infeasible, incomplete, or simply when only a single static sample is available.

A key advantage of Cavebear is its ability to anchor *in vitro* experiments to an *in vivo* developmental timeline. In this paper, we demonstrated the feasibility in the mouse embryoid body and human organoids. With the growing use of iPSC-derived cultures and organoids for modeling human diseases and testing drug-responses, we envision Cavebear will provide a quantitative framework for evaluating *in vitro* models and protocols, pinpointing development and maturation abnormalities across patients and disease conditions, as well as mapping *in vitro* perturbation effects to human physiology.

Beyond *in vitro* applications, Cavebear also provides unique opportunities in characterizing human disease progression. Due to the lack of time-series sampling at the disease sites and patient heterogeneity, characterizing where cells fall along a disease progression timeline remains challenging. Cavebear takes advantage of step-wise and gradual disease progression observations in inducted mouse cancer models or other model systems, to annotate cellular level patterns in patient samples. Here, we particularly showed how this is possible using a matched and unmatched cancer reference, highlighting the potential usage of references from different cancer types to find shared progression patterns. Our analysis was limited by available annotations, but we envision extensions of Cavebear where malignant cell clusters can more directly be linked to prognostic indicators.

Based on our experiences, several aspects inform reference selection. First, because the transfer learning framework relies on shared cell embeddings, Cavebear’s performance benefits most when there are shared cell types and overlapping time ranges between the reference and the query. Second, increasing the density of time labels of the reference improves pseudotime inference in the query, though a reference with several time labels already outperforms unsupervised methods. In addition, Cavebear achieves higher developmental pseudotime inference accuracy for cell types undergoing differentiation compared to cell types in relatively mature states, likely because transcriptional changes become less pronounced in the latter scenario. These observations highlight the importance of leveraging biologically appropriate references and suggest that increasingly comprehensive reference atlases will further improve the generalizability of pseudotime inference across species and biological conditions.

With the increasing effort of atlas scale measurements and perturbations, we envision Cavebear to benefit from more comprehensive time-series references to augment hypothesis generation where time annotations are missing. In addition, the current framework focuses on predicting pseudotime along a linear axis, and we expect incorporating branching trajectories in references could further refine continuous cell fate characterization. Finally, while we demonstrated Cavebear using single-cell RNA-seq data, as time-series from other single-cell modalities emerge, they may also benefit from Cavebear’s transfer-learning framework.

## Declarations

### Ethics approval and consent to participate

Ethics approval is not applicable for this study.

### Availability of data and materials

All datasets in this study are publicly available. Detailed information about the datasets is summarized in Table 1. The processed mouse reference datasets utilized in this study from Qiu et al. are available at https://omg.gs.washington.edu/jax/public/download.html as well as the raw datasets from NCBI Gene Expression Omnibus (GEO) using accession numbers GSE186069 and GSE228590. The processed mouse reference datasets utilized in this study from TOME can be accessed at https://tome.gs.washington.edu/ as well as the raw datasets from GEO using accession numbers GSE100597, GSE109071, GSE106587, GSE186069, and GSE186068, and from ArrayExpress (accession E-MTAB-6967). The zebrafish dataset is available at https://cole-trapnell-lab.github.io/zscape/downloads/ and GEO accession GSE202639. The embryoid body data can be accessed from GEO using the accession GSE184554. Human organoid raw data can be found at National Institutes of Mental Health Data Archive (NDA collection 4747), and processed data were provided directly by Jason Stein Lab at UNC-Chapel Hill. Colorectal cancer reference data is available at GSE260801, GSE296507, and human colorectal cancer patient scRNA-seq dataset is available at https://zenodo.org/records/10651059.

The Cavebear source code is available on Github at https://github.com/ranzhanglab/Cavebear.

## Acknowledgements

We would like to acknowledge Jason Stein and Esther Park for providing the processed organoid dataset and valuable suggestions. We would also like to thank Doudou Yu and Patrick Yu for providing feedback to the manuscript.

